# Acto3D: user- and budget-friendly software for multichannel high-resolution three-dimensional imaging

**DOI:** 10.1101/2023.08.18.553473

**Authors:** Naoki Takeshita, Shinichiro Sakaki, Rie Saba, Satoshi Inoue, Kosuke Nishikawa, Atsuko Ueyama, Kazuhiko Matsuo, Masaki Shigeta, Yoshiro Nakajima, Daisuke Kobayashi, Hideya Yamazaki, Kei Yamada, Tomoko Iehara, Kenta Yashiro

**Affiliations:** Division of Anatomy and Developmental Biology, Department of Anatomy, Graduate School of Medical Science, Kyoto Prefectural University of Medicine, Kyoto, Japan; Department of Pediatrics, Graduate School of Medical Science, Kyoto Prefectural University of Medicine, Kyoto, Japan; Department of Pediatrics, Graduate School of Medicine, The University of Tokyo, Tokyo, Japan; Department of Radiology, Graduate School of Medical Science, Kyoto Prefectural University of Medicine, Kyoto, Japan; Department of Pediatrics, Graduate School of Medicine, Osaka University, Osaka, Japan

**Author notes:** Correspondence: (N.T.) or (K. Yashiro).

## Abstract

Advances in fluorescence microscopy and tissue-clearing technology have revolutionized three-dimensional (3D) imaging of fluorescently labeled tissues, organs, and embryos. However, the complexity and high cost of existing software and computer solutions for such imaging limit its widespread adoption by researchers with limited resources. We here introduce Acto3D as a user- and budget-friendly, open-source computer software application designed to streamline the generation and observation of high-resolution 3D images of targets labeled with multiple fluorescent probes. Acto3D features an intuitive interface that simplifies the importation, visualization, and analysis of data sets, has an associated tool for annotation of vascular lumens, and incorporates multiple fluorescence channels for comprehensive imaging. Underpinned by an integrated graphics processing unit, Acto3D allows accurate image reconstruction and efficient data processing without the need for expensive high-performance computers. We validated the software by imaging mouse embryonic structures. Acto3D thus constitutes a cost-effective and efficient platform to support biological research.

---

With the advancement of tissue-clearing technology and fluorescence microscopy systems, it is now possible to label and identify deep structures of large tissues, organs, or organisms with the use of fluorescence probes^1^. However, it has remained a challenge to generate readily understandable three-dimensional (3D) images. Volume rendering has been a standard method of 3D imaging for visualization of internal to surface structures of an object based on definition of the intensity and opacity for each spatial coordinate^2^. In practice, 3D images obtained from ultrasonography, computed tomography (CT), or magnetic resonance imaging (MRI) via volume rendering provide much information in a visually and easily understandable manner^3^.

Volume rendering requires a powerful computer, more processing power and larger memory capacity as the space size increases. Specifically, images acquired by fluorescence microscopy often consist of a higher number of pixels in the XYZ dimension (incorporating multiple channels) compared with CT and MRI images in the clinics (Supplementary Fig. 1). Image size inevitably increases as the number of Z slices increases. Reconstruction of 3D images requires a graphics processing unit (GPU), which excels in fast parallel computations. If the GPU does not have sufficient dedicated memory, like that of the main processor of the system, the processing of such image data without downsizing or compression is a challenge. GPUs that possess a sufficiently large dedicated memory tend to be extremely expensive, however. In addition, volume rendering requires specialized software. Whereas many open-source software applications are available and offer highly advanced analysis functions^4–6^, they present difficulties with regard to loading multichannel raw data before compositing, to fine-tuning the display for each channel, and to dealing with loading times and processing speeds. On the other hand, there are commercially available software solutions, but these are invariably expensive and exceptionally demanding for high-spec hardware including dedicated GPUs (dGPUs). These factors impose a substantial barrier to adoption of such technology by researchers in their scientific activities. The ability to easily construct high-resolution 3D images at low cost would therefore greatly benefit the research community.

We here propose a pipeline of observation methods that can overcome these limitations. We have thus newly developed Acto3D—<u>A</u>daptive volume rendering software that allows users to freely adjust and customize <u>c</u>olor tones and transfer functions and provides for rich <u>o</u>bservation in <u>3D</u>—as an open-source volume rendering software application for multichannel fluorescence microscopic imaging. Acto3D takes advantage of the unified memory feature of Apple Silicon (Apple, Cupertino, CA, USA), which embodies the concept of fully deploying image data in the memory area accessible to a GPU. In Apple Silicon, both the central processing unit (CPU) and GPU access a unified memory with low latency^7^. This architecture can be regarded as incorporating a type of integrated GPU (iGPU) and allows the GPU to utilize a larger area of memory compared with that accessible to a typical dGPU installed in a personal computer (PC), resulting in extremely powerful graphics performance. This architectural advantage allows high-resolution volume rendering of microscopic images without the need for expensive workstations or the larger memory size designated for dGPUs. Furthermore, Acto3D provides a pipeline for extraction of spatial lumen structures and generation of mask images, as demonstrated by its ability to abstract 3D images of mouse embryonic great vessels. With validation of the usefulness and power of Acto3D in imaging of embryonic structures, we propose its adoption as a cost-effective and efficient tool.

## Results

### Overview of Acto3D volume rendering software

We developed Acto3D, using Swift and Metal Shading Language, as part of an open-source project and designed it to be a stand-alone application independent of external packages. This design concept eliminates the need for any specialized environment setup, making the application more accessible to a broader range of users. The primary focus of Acto3D is its use for 3D observation of fluorescence microscopic images. By allowing users to finely adjust opacity and color tones, Acto3D supports the free exploration of both surface and internal structures. It leverages the iGPU of Apple Silicon to provide a cost-effective platform that is optimized for Apple’s M-series chips. Of note, a high level of observation can be achieved even on a laptop machine (sup> (Fig. 1a and Supplementary Video 1). Depending on the image size, Acto3D can also be run on earlier series of Mac computers with a dGPU. A limitation is that it is exclusively an application for macOS.

**Fig. 1.**
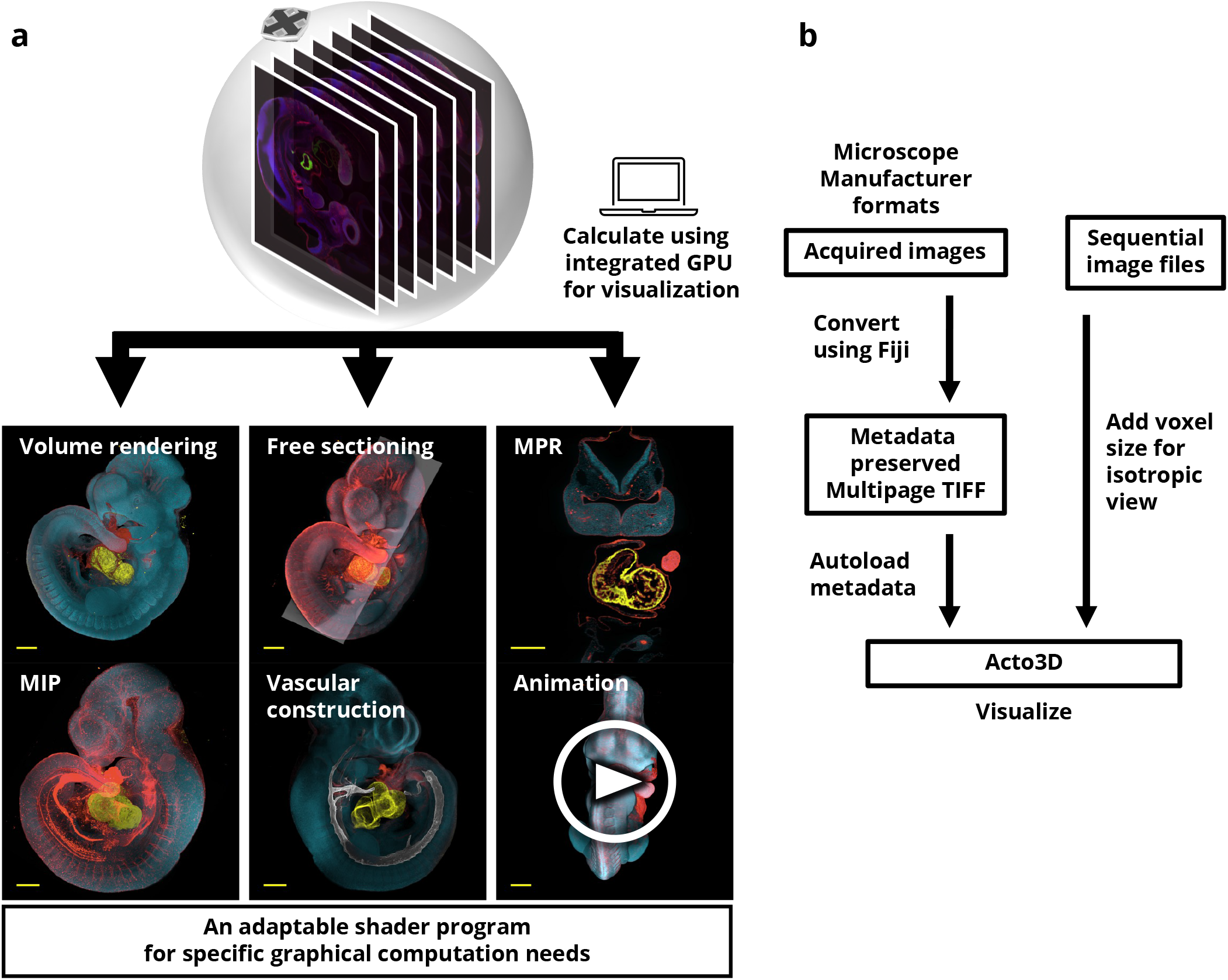
Overview of Acto3D. **a**, Stacks of the image are positioned in the correct spatial coordinates (considering the x, y, and z scales), and a virtual sphere that fully encompasses each stack is prepared. This virtual sphere can be observed from any angle, and the final pixel value is calculated according to the volume rendering equation. Various representations are possible with the use of different transfer functions. These calculations are performed by general-purpose computing with a GPU (GPGPU). Whereas Acto3D inherently allows for a variety of representations by default, it also anticipates more advanced usage, providing the flexibility to modify shaders partially or entirely as needed. MIP, maximum-intensity projection; MPR, multiplanar reconstruction. **b**, TIFF files saved by Fiji contain metadata such as voxel size and display ranges. Acto3D uses this information to construct 3D images with the correct dimensions. Simple loading of these TIFF files thus allows Acto3D to generate rich visualizations.

### Input image

Acto3D was designed to use z-stacks of multichannel images (up to four channels) captured by a fluorescence microscope. The XY dimension should be <2048 pixels (px), with up to 2048 slices in the Z dimension. Acto3D is able to handle sequential files in common image formats such as Tag Image File Format (TIFF), Portable Network Graphics (PNG), and Joint Photographic Experts Group (JPG). However, for utilization of the metadata of the raw image data acquired by the microscope system to present the object isotropically, ensuring consistent dimensions across all directions, the raw data should be exported as multipage TIFF by Fiji^8^ (Fig. 1b). Each channel of the image is clipped to 8 bits by Acto3D according to the specified display range. Depending on the number of channels, a 3D texture is generated: 8 bits for a single-channel image, 16 bits for a two-channel image, and 32 bits for a three- or four-channel image.

### Algorithm

In Acto3D, voxel data are sampled along the line of sight, and the resulting pixel values are composited with the rendering equation in common use^2, 9^ (Fig. 2a). Namely, with the back-to-front strategy:

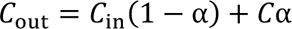

**Fig. 2.**
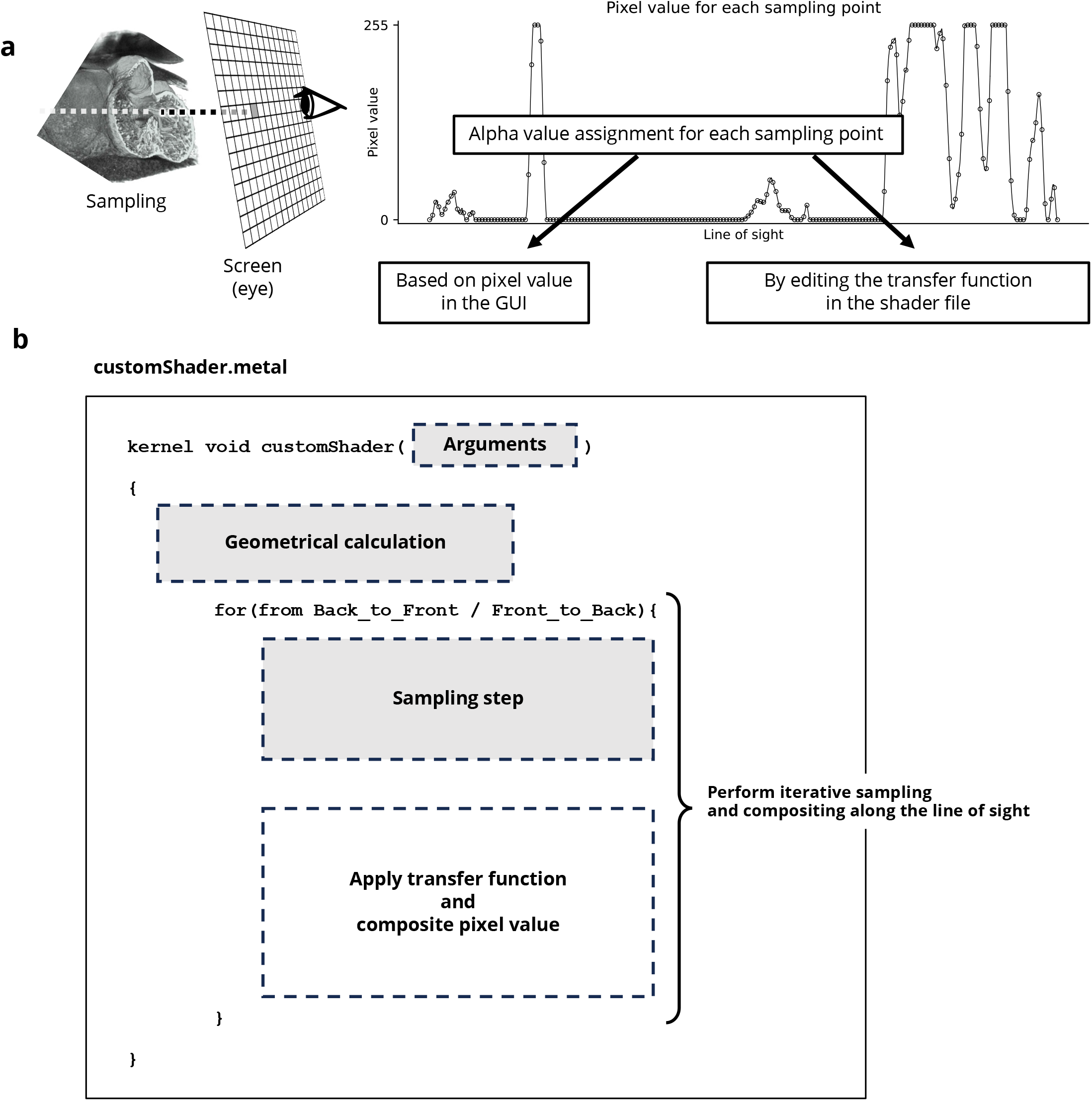
Overview of Acto3D operation and kernel shader. **a**, Pixel values are sampled along the line of sight during observation of the target object from any direction. Volume rendering requires opacity information for each sampling point. Primarily, the alpha value corresponding to the pixel value is determined by the graphical user interface (GUI), but it is also possible to edit the transfer function directly in the shader file, thereby providing more control for users with programming knowledge. **b**, Structure of the shader file. In Acto3D, the final pixel value is calculated by GPGPU, and the calculation process is explicitly defined in this shader file. A block for performing geometrical transformations is followed by a block that applies sampling and transfer functions in a front-to-back or back-to-front manner, with this process being iterated in a loop. If users wish to customize the kernel function, they have the flexibility not only to modify the stages of transfer function application and pixel value compositing but also to adjust across other blocks. Importantly, shaders can be compiled online, eliminating the need for compiling from the source code. It should be noted that users will need to write their modifications using Metal Shading Language, which is similar to C++.

With the front-to-back strategy, which has the advantage of being able to terminate calculations early, resulting in a lower computational cost^10–12^:

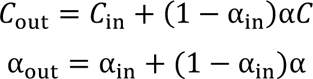

where *C*_in_ is accumulated intensity at the previous sampling point, *α*_in_ is accumulated opacity at the previous sampling point, *C* is the intensity of the current sampling point, *α* is the opacity of the current sampling point, and *C*_out_ and α_out_ are the output results of combining the previously accumulated sampling point and the present sampling point. All of these calculations are executed by compute kernels for general-purpose computing with a GPU (GPGPU) so as to clarify the processing steps for calculation.

The shader can be customized either partially or across the entire range to enhance display flexibility (Fig. 2b). Additions and modifications can be introduced without the need for recompiling the entire program from the source code. Shaders for multiplanar reconstruction (MPR) and maximum-intensity projection (MIP) were included as part of the initial set.

### Observation of whole mouse embryos

To validate Acto3D, we observed 3D models of embryonic day (E) 10.0 (32-somite stage) whole mouse embryos after tissue clearing with CUBIC-R+^13, 14^. Nuclear staining offered a comprehensive visualization of the entire shape of each embryo (Fig. 3a and Supplementary Fig. 2a). The surface structure was observed more readily as the overall opacity was increased, allowing somites and the pharyngeal pouches to be clearly distinguished via free rotation and sectioning (Fig. 3b,c). Conversely, as the opacity and intensity of the nuclear signals were reduced, internal structures became more readily visualized (Fig. 3d–f and Supplementary Fig. 2b,c). The detailed structure of the intersomitic vessel network was visible at increased magnification (Fig. 3e). Moreover, by customizing the transfer function to consider the gradient of the pixel value in the shader, we were able to highlight edges and thereby to identify the pharyngeal arch arteries (PAAs) and the dorsal aorta (Fig. 3f and Supplementary Fig. 2d). MPR images provided virtual slices from various perspectives that differed from the original imaging direction (Fig. 3g–i and Supplementary Fig. 3). Acto3D thus revealed readily comprehensible anatomic structures of organs and tissues in 3D.

**Fig. 3.**
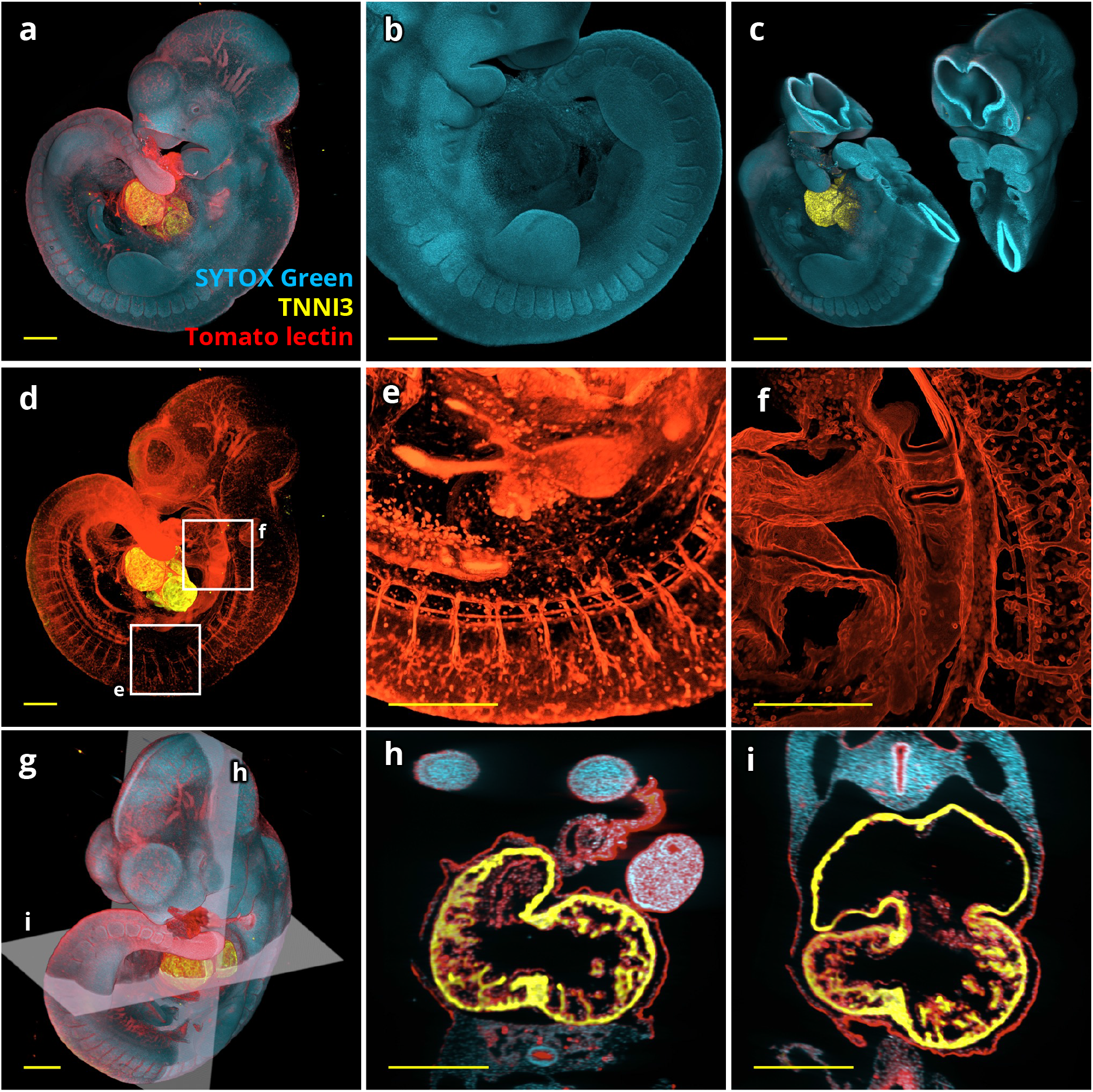
Observation of whole mouse embryos with diverse visualization modes. **a**, 3D whole-embryo image of a 32-somite mouse embryo stained with SYTOX Green to detect nuclei, with antibodies to cardiac troponin I (TNNI3), and with tomato lectin to detect blood vessels. The surface structure becomes visible as the opacity of all channels is increased. **b**, Image constructed solely from nuclear staining. The image is adjustable for observation from any angle, thereby facilitating the observation of somites. **c**, Cross-section of the pharyngeal pouch from two different angles. **d**, Image in which the opacity and intensity of the nuclear staining channel were reduced to emphasize vascular staining. **e**, Magnified view of the somite area (box e) in **d**. The intricate network of intersomitic vessels is apparent. **f**, The outlines of PAAs and the dorsal aorta are highlighted by adjustment of the transfer function to emphasize the gradient of surrounding area for box f in **d**. **g**–**i**, Acto3D is able to generate MPR images in any direction, irrespective of the imaging direction. The MPR images in **h** and **i** are for the sections in **g**. All scale bars, 500 µm.

### Observation of mouse embryonic heart development in 3D

We next observed development of the mouse embryonic heart^15–17^ (Fig. 4). At E9.5, the presumptive left and right ventricles were discernible (Fig. 4a) and trabeculae were well developed in the ventricles (Fig. 4b,c). At E10.5, the onset of formation of the muscular ventricular and primary atrial septa was apparent^15, 18^ (Fig. 4e,g). The atrioventricular valves still retained the superior and inferior atrioventricular cushions, connecting only to the left ventricle (Fig. 4e,f). At E11.5 or later, the rightward shift of the atrioventricular canal responsible for its connection to both ventricles was seen (Fig. 4i,m). The primary atrial septum had further developed, and a secondary foramen had begun to form (Fig. 4i,j). The outflow tract still originated only from the right ventricle and was not separated into pulmonary and aortic trunks (Fig. 4k). By E12.5, the atrioventricular cushions were separated into the presumptive mitral and tricuspid valves (Fig. 4m). The outflow tract was separated into pulmonary and aortic trunks, and its stem was shifted leftward, leading to positioning of the aortic valve over the muscular ventricular septum (Fig. 4n). The opening of the pulmonary vein was observed near the dorsal rim of the atrial septum on the left atrial side (Fig. 4n). Presumptive papillary muscles were also clearly identified at this stage (Fig. 4o), which shows the utility of Acto3D because this had not been confirmed by a conventional method to date^16^. All these observations were thus consistent with those of previous studies^15–18^.

**Fig. 4.**
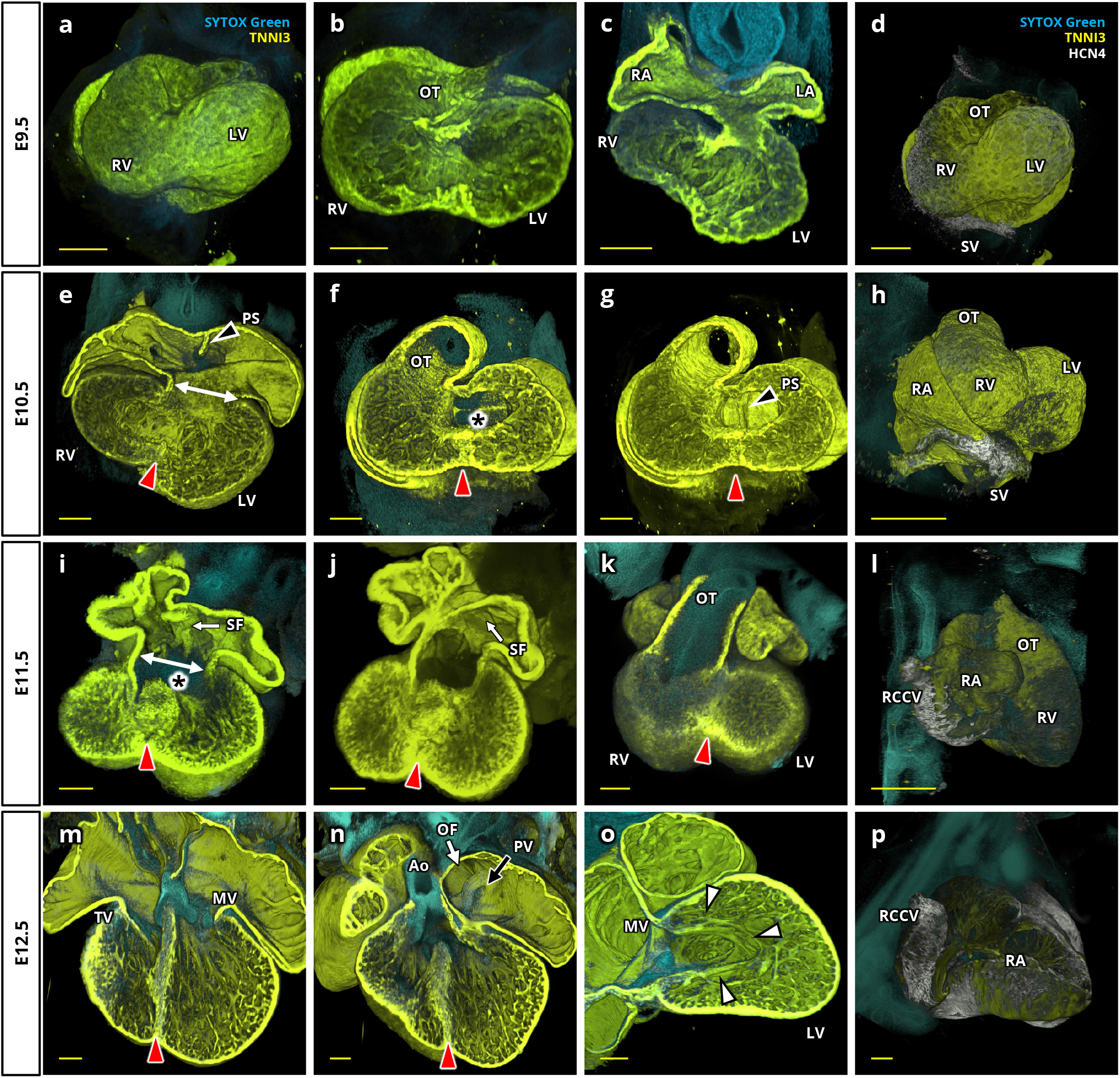
Morphological changes during development of the mouse embryonic heart. **a**–**c**, The heart of an E9.5 embryo stained with SYTOX Green and antibodies to TNNI3. An overall 3D image of the heart is shown in **a**, with coronal and transverse sections being presented in **b** and **c**, respectively. **e**–**g**, Transverse and coronal sections of the heart at E10.5 are shown in **e** and in **f** and **g**, respectively. The atrioventricular canal (white two-way arrow in **e**) connects only to the left ventricle (LV), whereas the outflow tract (OT) connects only to the right ventricle (RV). Black arrowheads with a white border indicate the primary atrial septum (PS). **i**–**k**, The heart of an E11.5 embryo. The atrioventricular canal is seen to communicate with both ventricles (**i**), and a slitlike secondary foramen (SF), indicated by the white one-way arrow, has formed dorsally. The OT still originates entirely from the RV (**k**). **m**–**o**, The heart at E12.5. The mitral valve (MV) and tricuspid valve (TV) have formed (**m**). In **n**, the aorta (Ao) is positioned over the ventricular septum, and the opening of the pulmonary vein (PV, indicated by the black arrow with a white border) is clearly observed near the oval foramen (OF, indicated by the white arrow with a black border). A papillary muscle (white arrowheads with a black border) was identified (**o**). **d**, **h**, **l**, **p**, HCN4 distribution at E9.5, E10.5, E11.5, and E12.5, respectively. HCN4 was initially found in the cardiac region where the early sinoatrial node develops, but its expression gradually extended to encompass the common cardinal veins. All scale bars, 200 µm. LA, left atrium; RA, right atrium; SV, sinus venosus; RCCV, right common cardinal vein. Asterisks (*) indicate the atrioventricular cushion, and red arrowheads indicate the muscular ventricular septum.

HCN4, a marker of the early electrical conduction system including the sinoatrial node^19–21^, was initially localized to the sinus venosus (the inflow tract) at E9.5 and gradually extended to the region where the right and left common cardinal veins enter the sinus venosus (Fig. 4d,h,l,p and Supplementary Fig. 4). The intensity of HCN4 subsequently increased in the common cardinal veins in a left-right–asymmetric manner. Consistent with previous observations^21^, the common cardinal vein was also found to be positive for cardiac troponin I (TNNI3) (Supplementary Fig. 4 and Supplementary Video 2).

These various observations thus indicated that Acto3D facilitates the observation of structures of interest by rendering them more discernible. More importantly, it allowed sectioning of the heart at various angles and thereby provided a vivid depiction of the morphological changes in the internal structure of the embryonic heart.

### Abstraction of great vessels via spatial clustering

During embryogenesis, the morphology of PAAs changes markedly through remodeling^22, 23^. We next aimed to observe PAAs through challenging reconstruction from mask images obtained by filling in the vascular lumen edged with fluorescently labeled endothelium. Imaging by fluorescence microscopy usually needs to overcome two key problems: variation in staining intensity among specimens, and the attenuation of emission light intensity that is either caused directly by the presence of material between the target object and the objective lens or due to a reduction in excitation light intensity reaching the target during its passage through intervening tissue^24^. It is therefore not feasible to automatically recognize the vascular lumen by setting a specific threshold for intensity values and filling in the entire lumen.

To address this issue, we iteratively applied one-dimensional k-means clustering^25^ spatially to single slice images (Fig. 5a). We first prepared data by removing noise through appropriate preprocessing and cropped the local vascular area of interest. We next applied the k-means++ algorithm^26^ to the first slice of this area in order to obtain the initial centers, which were then used to obtain initial centroids of cluster classification via standard k-means clustering. Given that the selection of initial centers by k-means++ relies on a weighted probability distribution, the calculation results differed each time. If unintended cluster classification occurred, we either reran the calculation or changed the number of clusters. If imaging was performed with thin slices, adjacent slices had structurally similar images and similar intensities as revealed by the Structural Similarity Index Measure^27^ (Supplementary Fig. 5a). As expected, adjacent images had similar classifications after specifying the obtained centroids in one slice as the initial centers for the next slice (Fig. 5b). The continuous slices obtained in this manner showed a smoother transition with reduced computational costs compared with the application of k-means++ to each independent slice one by one (Supplementary Fig. 5b–e). Importantly, the vascular structure constructed from the mask images obtained by this iterative method represented well the structure of the vascular lumen with high resolution (Fig. 5c). Using this method, we were able to construct a 3D model of PAAs with the information of their surrounding tissues including the embryonic pharyngeal pouches (Fig. 6a–c and Supplementary Video 3).

**Figure 5.**
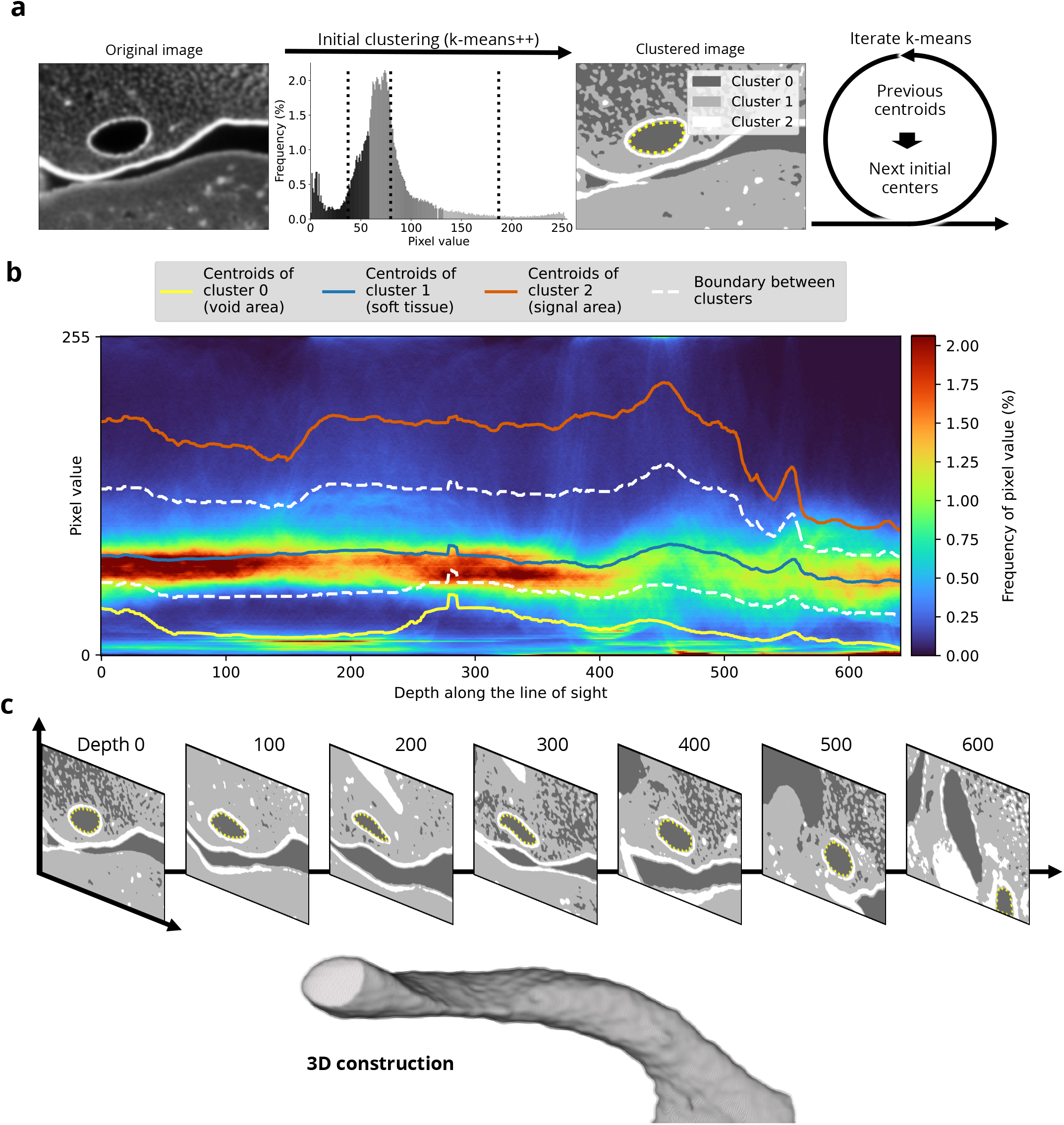
Overview of spatial vascular tracing. **a**, Images of the area of interest are clipped and subjected to clustering with the k-means++ algorithm for selection of the initial center points. The centroids resulting from this step are then used as the initial center points for clustering in the next slice. Note that random numbers are used only in selection of the initial center points in the first slice. The dashed lines in the histogram represent the calculated centroids for the initial slice. In the clustered image, the target vascular lumen is indicated by the yellow dashed line. **b**, With the use of the centroids of the previous slice as the initial centers for the next slice, clustering can be performed down to the deepest level while maintaining the trend of the first clustering. **c**, A 3D reconstruction of a blood vessel that was generated from the mask images obtained by this iterative process.

**Figure 6.**
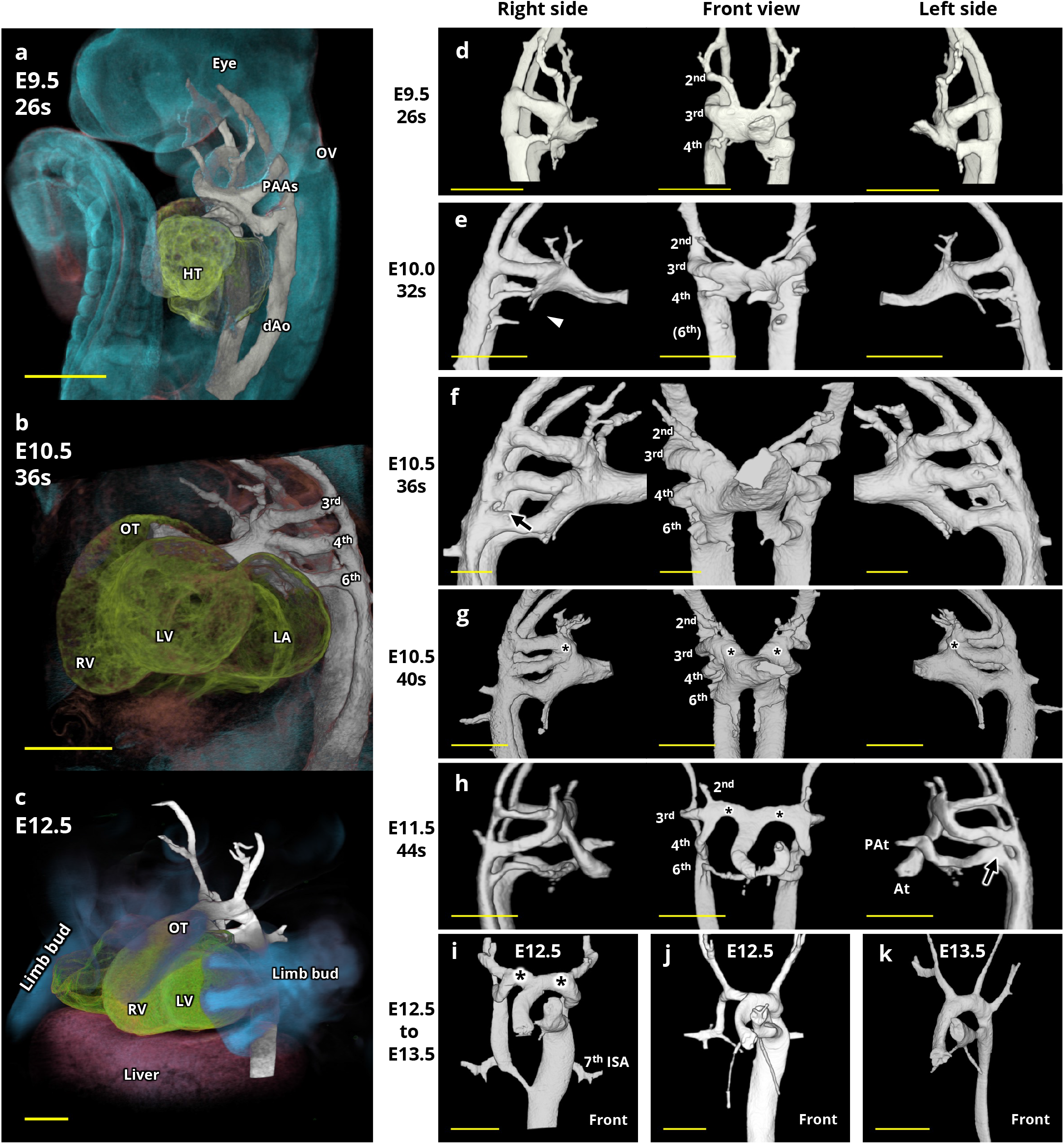
Remodeling of PAAs. **a**–**c**, Left frontal views of PAAs and surrounding tissues in mouse embryos at E9.5, E10.5, and E12.5, respectively. The embryos were stained with SYTOX Green (blue), antibodies to TNNI3 (green), and tomato lectin (red). Constructed PAAs are shown in white. dAo, dorsal aorta; HT, heart; OV, otic vesicle; s, somite; OT, outflow tract; RV, right ventricle; LV, left ventricle; LA, left atrium; 3rd, 4th, and 6th indicate the third, fourth, and sixth PAAs. Scale bars, 500 µm. **d**–**k**, 3D reconstruction models depicting the morphology of PAAs viewed from the right lateral, front, and left lateral sides and at the indicated developmental stages. At the 32-somite stage, the intermediate portion of the sixth PAA was not constructed (white arrowhead), although the endothelium continuously connected the aortic sac and dorsal aorta (Supplementary Fig. 6). Some embryos possessed a dorsal collateral connecting the fourth and sixth PAAs (black arrow in **f** and **h**). At, aortic trunk; PAt, pulmonary artery trunk; 7th ISA, seventh intersegmental artery. Asterisks (*) indicate the aortic sac horn. Scale bars, 500 µm.

### Observation of high-resolution 3D models of PAAs

We observed the chronological 3D model of PAA development^22, 23^ (Fig. 6d–k). At E9.5 (26-somite stage), the second, third, and fourth PAAs were seen (Fig. 6d). By E10.0 (32-somite stage), the second PAA had lost its connection with the dorsal aorta, and formation of the sixth PAA appeared to be under way in the 3D model (Fig. 6e). However, observation of the acquired images used for construction of this 3D model revealed that the endothelium of the presumptive sixth PAA, while lacking a visible lumen, was continuously connected between the aortic sac and dorsal aorta (Supplementary Fig. 6). At E10.5 (36-somite stage), formation of the sixth PAA was complete (Fig. 6f). At the 40-somite stage, an aortic sac horn had formed as a common duct from the aortic sac to the third and fourth PAAs, resulting in a slight anterior shift in the positions of these arteries away from the heart (Fig. 6g). At E11.5 (44-somite stage), the aortic sac had separated into the aortic and pulmonary trunks, and the right sixth PAA had started to regress. The base of the outflow tract underwent a rotation, resulting in movement of the aortic trunk posteriorly and relocation of the pulmonary trunk anteriorly (Fig. 6h). Around E12.5, the right dorsal aorta of the periphery started to regress below the level of the seventh intersegmental artery (Fig. 6i,j). At E13.5, this portion of the aorta had completely regressed and the morphology of the great artery system was similar to that of an adult (Fig. 6k). All our observations of 3D images of PAAs were thus consistent with previous findings^22, 23, 28^.

In some embryos, we were able to confirm the presence of dorsal collateral vessels between the fourth and sixth PAAs (Fig. 6f,h). It is proposed that persistent fifth aortic arch is not a remnant of the fifth pharyngeal arch artery itself, but a morphological abnormality caused by these collaterals^28–31^.

## Discussion

We here describe development of software, Acto3D, that allows easy 3D reconstruction for flexible observations from multichannel z-stack fluorescence images. Acto3D is a robust platform designed to allow fine-tuning of opacity and highlighting of target structures, providing sophisticated and readily understandable 3D anatomic images for viewing from multiple perspectives. We have validated the benefit of this software for detailed characterization of anatomic structures in 3D. It thus provided sophisticated 3D models both of the mouse embryonic heart and, with the support of an algorithm for tracing vessel lumens, of PAA development. This powerful tool can theoretically be applied not only to embryonic cardiovascular structures but to other targets with appropriate probe labeling and tissue-clearing technology. With its ability to observe 3D morphology as well as to visualize internal structures, Acto3D should prove to be of great benefit in various life science fields.

Previous studies have presented 3D interpretations of the mouse embryonic heart obtained with the use of various techniques including episcopic microscopy^15–18^ and micro-CT^32^. These studies focused primarily on depicting general morphology without observation of specific structures derived from specific cell lineages. 3D models generated by Acto3D from acquired fluorescence images allow general morphological observations based on nuclear staining only. More importantly, it is possible to study a target structure or the distribution of target cells revealed by specific immunostaining within the general anatomic structure with the use of multichannel 3D images. This task can be smoothly handled even on a laptop computer (Supplementary Video 1). Theoretically, even a low-magnification lens, such as a 5× magnification objective, should yield a pixel resolution of ∼1 to 2 µm, which we found sufficient for visualization of individual trabeculae inside the embryonic heart ventricles.

Compared with previous studies of the 3D morphology of PAAs^22, 28^, the application of Acto3D to obtain 3D models of these arteries had several advantages including technical simplicity, low cost, high resolution, the ability to observe cross-sections and from different viewing directions, and the provision of information on surrounding structures. One limitation was that vascular lumens could not be observed if the vessels were not patent or were too thin. The use of confocal microscopy instead of light sheet microscopy may be more suitable for further detailed local observations as previously suggested^33^.

In general, microscopy images have a lower resolution in the Z direction than in the XY direction. Even if the voxel size is isotropic, it is therefore impossible for all resolutions to be equal in space. Depending on the construction method for 3D images, it may be necessary to adjust the arrangement of specimens and align the plane of the light sheet in order to achieve the maximum resolution. However, with the use of Acto3D, we found that the influence of specimen orientation was small in MPR imaging and also negligible in the case of 3D observation. Positioning of the specimen in the desired orientation for observation of small structures by light sheet microscopy can be challenging. However, with the use of Acto3D for 3D observation, the impact of placement direction on image quality can be minimized (Supplementary Fig. 3).

The use of Apple Silicon played a substantial role in the development of our pipeline, largely as a result of its good fit for our dual objectives of deploying the entirety of image data in GPU-accessible memory and allowing meticulous observations of fine details at any desired time. However, the consequence of this reliance on Apple Silicon is that Acto3D is currently limited to macOS. The use of an external GPU to achieve high performance even with a laptop machine did not align with our objectives because of the more complex configuration required. There is the potential for cross-platform applicability of Acto3D in the future if similar high-performance iGPUs are developed or for PCs with large-capacity dGPUs.

## Methods

### Mice

ICR mice were obtained from Japan SLC. All animal procedures were approved and performed according to guidelines specified by the Animal Experimentation Committee of Kyoto Prefectural University of Medicine (license nos. M2022-172 and M2022-180). The morning of the day on which a vaginal plug was detected after breeding was determined as E0.5. Embryos were dissected at times from E9.5 to E13.5. The impact of blood autofluorescence was minimized by flushing out of the vessels of an embryo through injection of 4% paraformaldehyde in phosphate-buffered saline (PBS) with the use of a glass needle connected to a 1-ml syringe (Terumo) and an Injection Holder (IM-H1, Narishige Scientific Instrument Lab). The glass needles were prepared with the use of a puller (PC-10, Narishige Scientific Instrument Lab) set at 52°C from borosilicate glass capillaries with outer and inner diameters of 1.0 and 0.58 mm, respectively (GC100F-10, Harvard Apparatus). For observation of the heart, the abdomen of the embryo was punctured and 4% paraformaldehyde in PBS was gradually injected to identify the appropriate injection site for flushing out of blood cells from the heart and great vessels. After visual confirmation with a stereomicroscope that blood cells had been adequately removed from the ventricles, any unwanted tissue was trimmed. For 3D reconstruction of vessels, the ventricles were punctured and blood cells were flushed out by manual injection until no further such cells flowed out from the umbilical cord. In both cases, after perfusion with 4% paraformaldehyde, the specimens were incubated overnight at 4°C in the same fixative.

### Immunofluorescence analysis and tissue clearing

Immunostaining was performed as previously described^13^, with some modifications. After fixation, specimens were washed three times for 30 min with PBS at room temperature. Delipidation was performed with 10% *N*-butyldiethanolamine and 10% Triton X-100 in deionized distilled water (DDW) (CUBIC-L) at 37°C for 24 to 48 h, until the CUBIC-L no longer changed color. The samples were then washed three times for 2 h with PBS. For nuclear staining, they were incubated overnight at 37°C with SYTOX Green (S7020, Thermo Fisher Scientific) at a dilution of 1:2500 in a solution of 10% Triton X-100, 5% *N*,*N*,*N’*,*N’*-tetrakis(2-hydroxypropyl)ethylenediamine (Quadrol #T0781, TCI,), 10% urea (#35904-45, Nacalai Tesque), and 500 mM NaCl in DDW (Sca*l*eCUBIC-1A with 500 mM NaCl). They were then washed three times for 2 h with 10 mM HEPES-NaOH (pH 7.5). For prevention of nonspecific binding of probes, the specimens were incubated for 90 min at 32°C in a solution containing 10 mM HEPES-NaOH (pH 7.5), 10% Triton X-100, 200 mM NaCl, 0.5% casein (#030-01505, Fujifilm Wako), 2.5% *N*,*N*,*N’*,*N’*-tetrakis(2-hydroxypropyl)ethylenediamine, and 1 M urea (HEPES-TSC buffer with additives). They were then incubated for 3 days at 32°C with an appropriate combination of primary antibodies to TNNI3 (ab56357, Abcam; diluted 1:200 to a concentration of 5 µg/ml), DyLight 594–conjugated tomato lectin (DL1177, Vector Laboratories; diluted 1:150), and/or antibodies to HCN4 (AB5808, Millipore; diluted 1:250) in HEPES-TSC buffer with additives. The samples were washed first for 2 h at 32°C with 0.1 M phosphate buffer at pH 7.5 (PB) containing 10% Triton X-100 and then three times for 2 h at room temperature with PB. After incubation for 90 min at 32°C with HEPES-TSC buffer with additives, the samples were exposed for 2 days at 32°C to secondary antibodies diluted in HEPES-TSC buffer with additives including an appropriate combination of Alexa Fluor 555–conjugated donkey antibodies to rabbit immunoglobulin G (#A-31572, Invitrogen; diluted 1:250) for HCN4 and/or Alexa Fluor 633–conjugated donkey antibodies to goat immunoglobulin G (#A-21082, Invitrogen; diluted 1:200) for TNNI3. Samples were washed once for 2 h at 32°C with PB containing 10% Triton X-100 and then three times for 2 h at room temperature with PB. The stained embryos were then incubated with 1% formaldehyde (#16223-55, Nacalai Tesque) in PB first for 60 min at 4°C and then for 45 min at 37°C. They were washed three times at room temperature with PB. Refractive index matching was performed by incubation of the samples first overnight in a solution of 45% 2,3-Dimethyl-1-phenyl-5-pyrazolone (Antipyrine, #D1876, TCI) and 30% N-Methylnicotinamide (#M0374, TCI) in DDW (CUBIC-R+)^13, 14^ diluted to 0.5× with DDW, and then for at least 16 h in CUBIC-R+ at room temperature.

### Image acquisition by light sheet microscopy

Images were acquired with a Zeiss Lightsheet 7 fluorescence microscope. An EC Plan-NEOFLURA 5×/0.16 foc objective lens was used for detection, and an LSFM 5×/0.1 foc objective lens for illumination. The laser setup was applied only on the left side. Each specimen was embedded in a glass tube with an inner diameter of 2.2 mm (#701910, BRAND) or 1.5 mm (#701908, BRAND). If an appropriate-sized glass tube was not available, the specimen was embedded in 2% agarose (#02468-95, Nacalai Tesque) in CUBIC-R+. With regard to acquisition parameters, the zoom was set between 0.36 and 1.16 to ensure that the area of interest fitted within a frame of 1920 by 1920 pixels; the center thickness was adjusted to maintain an optimized Z interval/XY pixel ratio of <2; the laser output for each wavelength was set within the range of 10% to 50% so as to avoid color saturation; and the exposure time was fixed at 99.87 ms. In general, image stacks consisted of 600 to 1200 z-slices and were saved in 16-bit depth.

### Volume visualization with Acto3D

For volume visualization with the use of Acto3D, the development environment consisted of an Apple MacBook Pro (model MacBookPro18,4) equipped with an Apple M1 Max processor featuring 64 GB of memory and running on OS version 12.5.1. In addition, an Apple M1 Pro processor with 32 GB of memory, and Apple M1 processor with 16 GB of memory, and an Apple M2 processor with 16 GB of memory were used for verification. Although not officially declared by Apple Inc., the iGPU had limitations in securing memory capacity all at once, allowing for approximately half of the total memory to be allocated. Specifically, for the M1 Max with 64 GB of memory, the maximum buffer size was 36 GB; for the M1 Pro with 32 GB of memory, it was 16 GB; and for the M1 with 16 GB of memory, it was 8 GB. It was therefore essential to ensure that the size of each image adhered to these specific constraints.

We first loaded the Zeiss czi format using Fiji^8^ (version 2.3.0/1.53s). The import options were set to Grayscale color mode, with Virtual Stack turned on and Split Channel turned off. We then converted the file to multipage TIFF format using the [File] → [Save As] → [Tiff…] function. At this stage, crucial metadata, such as display ranges and voxel size, were stored in the TIFF file. To achieve 3D visualization, we loaded this TIFF file into Acto3D according to the following procedure: Acto3D → [Open Images] → [Open ImageJ / Fiji TIFF]. We then set the display range for each channel through [Image Option…]. Given that pixel values in fluorescence microscopy can vary as a result of factors such as laser output, staining conditions, and XYZ position, there are no universally applicable recommended parameters. Nonetheless, we generally adjusted the display range to avoid intensity saturation and to minimize background noise. The [View in 3D] button was used to execute the volume visualization. For each channel, the opacity level and intensity could be adjusted through the graphical user interface (GUI), as detailed in Supplementary Figure 2. Of note, the final image resulting from volume rendering may exhibit variation due to the number of overlapping images, even with the same settings. No absolute parameters can therefore be considered universally perfect for all cases. However, we generally achieved good rendering images by setting the opacity to 0 near brightness level 0, assigning low opacity to low-intensity areas, and adopting medium to high opacity for high-intensity areas. This configuration served as a basis for fine-tuning the opacity for each brightness value in each channel, allowing us to obtain optimal images. Several settings can be found in our GitHub repository: https://github.com/Acto3D/Acto3D.

### 3D reconstruction of PAAs with Acto3D

Images of specimens labeled with tomato lectin were loaded into Acto3D with the same approach as in the previous section. The display range was set high enough to capture the signal value of endothelial cells of the blood vessels. A dedicated screen was opened through [Segment] → [3D Segment], where we selected the channel that imaged tomato lectin, and [Preprocess] was executed. This step involved application of a Gaussian blur to the 3D data with a kernel size of 7 × 7 × 7. From the displayed MPR images, we selected the blood vessels constituting the PAAs and chose the first and last slices for construction using [Set]. We next set the number of k-means++ clusters (usually, 3 or 4) and performed the initial clustering with the reload button. Given that k-means++ clustering relies on weighted probability, the calculation results may vary with each execution. After the necessary cluster classification was obtained, we selected the cluster area representing the blood vessel lumen and pressed [Start] or [Resume] button to track the same cluster according to the aforementioned algorithm. This process was repeated for multiple blood vessels constituting the PAAs, and their data were finally merged and incorporated into the visualization by pressing [Apply To Main]. Detailed instructions for these operations can also be found in our GitHub repository: https://github.com/Acto3D/Acto3D.

### Code availability

The source code, compiled binaries, and comprehensive documentation for the application, including guidelines on generating custom shaders, can be accessed through the GitHub repository (https://github.com/Acto3D/Acto3D).

## Acknowledgements

This work was supported by JSPS (Japan Society for the Promotion of Science) KAKENHI Grant-in-Aid for Scientific Research (B) (no. JP23H02878), JSPS KAKENHI grant (ABiS) (no. JP22H04926), and grants from Kawano Masanori Memorial Public Interest Incorporated Foundation for Promotion of Pediatrics and Miyata Foundation Bounty for Pediatric Cardiovascular Research to K. Yashiro; and by JSPS Grant-in-Aid for JSPS Fellows (DC2) (no. JP22J12994) to N.T. . We thank the Kyoto University Live Imaging Center (KULIC) for the acquisition of imaging data with the Zeiss Lightsheet 7 microscope as well as Michiyuki Mastuda and Etsuo Susaki for technical advice and helpful discussion.

## Author contributions

K. Yashiro conceived and designed the study. K. Yashiro and N.T. planned the experiments. N.T. performed the experiments as well as designed and constructed Acto3D. K.N., A.U., S.I., S.S., R.S., K.M., M.S., Y.N., D.K., H.Y., K. Yamada, and T.I. contributed to the analyses and interpretation of data. K. Yashiro and N.T. wrote the manuscript. All authors discussed the results and commented on the manuscript.

## Competing interests

The authors declare no competing interests.

## Supplementary Figure Legends

**Supplementary Fig. 1. Comparison of data size between medical imaging and fluorescence microscopy.** Data size per slice is compared for typical MRI (256 by 256 px), typical CT (512 by 512 px), ultra–high-resolution CT (2048 by 2048 px), and fluorescence microscopy (for example, 1920 by 1920 px for four-channel imaging with Zeiss Lightsheet 7). The right axis indicates the image data size when 500, 750, or 1000 slices are imaged.

**Supplementary Fig. 2. Adjustment of visualization. a**, Acto3D provides a panel within the software screen that allows the user to adjust color tones and the transfer function (white box). In addition to color tone, the user is thus able to adjust the intensity of each channel and the opacity corresponding to pixel values. The intensity setting multiplies the pixel value at each sampling point in each channel, so that increasing it makes the image brighter. Opacity is lowered for regions with low pixel values and set higher for those with high pixel values. In volume rendering, even very low opacities can make it difficult to see through the object if overlaid in large amounts; opacity is therefore squared and internally adjusted by default (this can be changed). The mouse embryo showcased here corresponds to the stained embryos depicted in Figure 3. **b**, By increasing the intensity of the nuclear staining image, the surface of the entire embryo can be viewed. **c**, By reducing the intensity and opacity of the nuclear staining image, the internal structure can be viewed. **d**, For emphasis of edges, the norm of the difference in pixel value of surrounding points is used instead of the pixel value at the sampling point. It becomes possible to use this flexible transfer function by construction of a custom shader.

**Supplementary Fig. 3. Construction and visualization of 3D structures and MPR images from different illumination directions and sample orientations.** For 3D reconstructions, Acto3D eliminates the need to precisely arrange the specimen and to align the light sheet plane in order to achieve maximum resolution. Comparison of data obtained from the same E10.0 mouse embryo stained as in Figure 3 by anteroposterior scanning in the coronal plane (**a**) with those obtained by lateral scanning in the sagittal plane (**b**) allowed the acquisition of XZ or YZ plane images that were similar in resolution to the theoretically highest-resolution plane image in the XY plane. MPR images are shown in the right panels of **a** and **b**. All scale bars, 500 µm.

**Supplementary Fig. 4. 3D observation of HCN4 expression**. The images depict views from the dorsal side of the heart of mouse embryos at E9.5, E10.5, E11.5, and E12.5, including MPR images from the same viewport showing the sinus venosus (SV) and common cardinal veins (CCVs). The embryos were stained with SYTOX Green and with antibodies to HCN4 and to TNNI3. The HCN4 signal was initially localized near the SV on the ventral side of the heart and gradually advanced upward toward the CCVs. The expression of HCN4 became left-right asymmetric in association with the left-right asymmetric remodeling of the cardinal veins. HCN4 expression overlapped with that of TNNI3. All scale bars, 500 μm. LA, left atrium; RA, right atrium; RHSV, right horn of SV; LCCV, left common cardinal vein; RCCV, right common cardinal vein.

**Supplementary Fig. 5. Evaluation of the efficiency of iterative and one-by-one k-means++ approaches. a**, The similarity between adjacent slices along the line-of-sight direction is plotted as the Structural Similarity Index Measure (SSIM), which reflects similarity in structure and intensity. The high SSIM values of >0.98 indicate similarity in intensity and structure between adjacent slices. **b**, If k-means++ is applied independently to each cross-section—for instance, by setting the total number of clusters to 3—random numbers are assigned to each cluster in each cross-section. Note that cluster centroids of high, medium, and low pixel values are not consistently allocated to each cluster. **c**, When k-means++ is applied independently to each cross-section, clusters corresponding to the vascular lumen are assigned random numbers for each cross-section. **d**, The computed k-means++ clusters identified for each cross-section one-by-one in **b** were sorted to match those that should be classified into the same cluster on the image. Application of k-means++ independently to each cross-section one-by-one therefore entails additional computational costs. Note the lack of smoothness in the graph representing the pixel values of the cluster centroids in comparison with the results obtained by the iterative k-means++ method (Fig. 5b). **e**, The clustered grayscale images obtained by the iterative (Fig. 5c) or one-by-one (**c**) k-means++ approaches were subjected to SSIM calculation. The absolute difference in SSIM was computed as |SSIM_*n* + 1_ − SSIM_*n*_|, where SSIM_*n*_ denote the SSIM values calculated from the *n*th and (*n* + 1)th clustered images, respectively. The results reveal that neighboring images are more similar with the iterative method (with the absolute difference in SSIM being close to 0) compared with the one-by-one k-means++ approach.

**Supplementary Fig. 6. Issues in constructing 3D models of extremely narrow blood vessels. a**, A 3D reconstructed image of PAAs in a mouse embryo at E10.0 (32-somite stage). The specimen is the same as that in Figure 6e. **b**,**c**, MPR views of the same embryo as in **a** are shown as sagittal (**b**) and frontal (**c**) sections. In the 3D model (**a**), the sixth PAA did not appear continuous between the aortic sac and dorsal aorta. However, in the MPR views of the region corresponding to that where the sixth PAA was expected to be identified in **a** (dashed black square), a continuous line of endothelium (black arrowheads with a white border) was observed between the aortic sac and dorsal aorta, suggesting that the artery was not constructed as a result of it being too thin or not possessing a lumen. The endothelial structure of the pulmonary artery (white arrow with a black border) was observed, but due to its extreme thinness, it could not be constructed. Scale bars, 200 µm.

**Supplementary Video 1. Operation overview of Acto3D.** Acto3D enables effortless manipulation of opacity, color tones, and transfer functions in multi-channel fluorescent images, facilitating comprehensive 3D visualization with optimal processing speed even on a laptop. The shaders governing rendering techniques and transfer functions, including multiplanar reconstruction (MPR), can be instantaneously toggled, enhancing adaptability during visualization.

**Supplementary Video 2. HCN4 expression in 3D space for a mouse embryo at E11.5.** The embryo was stained with SYTOX Green (blue) and with antibodies to TNNI3 (yellow-green) and to HCN4 (white). The atrium and common cardinal veins are positioned so as to sandwich the trachea in a V-shape. Pronounced expression of HCN4 was observed on both sides of the common cardinal veins and the right atrium. Scale bar, 500 μm.

**Supplementary Video 3. Segmentation of PAAs.** Initially, the region of interest is cropped, specifying the start and end slices. The number of clusters is determined empirically to extract the desired structures, first by determining the initial centers using k-means++. Subsequently, by specifying the cluster centroids of the previous slice as the centers for the next slice and iteratively applying k-means automatically, the desired structures can be efficiently extracted.

## Notes

https://github.com/Acto3D/Acto3D

### Competing Interest Statement

The authors have declared no competing interest.

## References

1. Kolesova, H., Olejnickova, V., Kvasilova, A., Gregorovicova, M. & Sedmera, D. Tissue clearing and imaging methods for cardiovascular development. iScience 24, 102387 (2021).

2. Levoy, M. Display of surfaces from volume data. IEEE Computer Graphics and Applications 8, 29–37 (1988).

3. Spanaki, A. et al. 3D Approaches in Complex CHD: Where Are We? Funny Printing and Beautiful Images, or a Useful Tool? J Cardiovasc Dev Dis 9 (2022).

4. Peng, H., Ruan, Z., Long, F., Simpson, J.H. & Myers, E.W. V3D enables real-time 3D visualization and quantitative analysis of large-scale biological image data sets. Nat Biotechnol 28, 348–353 (2010).

5. Peng, H., Bria, A., Zhou, Z., Iannello, G. & Long, F. Extensible visualization and analysis for multidimensional images using Vaa3D. Nat Protoc 9, 193–208 (2014).

6. Schmid, B., Schindelin, J., Cardona, A., Longair, M. & Heisenberg, M. A high-level 3D visualization API for Java and ImageJ. BMC Bioinformatics 11, 274 (2010).

7. Kenyon, C. & Capano, C. in 2022 IEEE High Performance Extreme Computing Conference (HPEC) 1–10 (2022).

8. Schindelin, J., et al. Fiji: an open-source platform for biological-image analysis. Nat Methods 9, 676-682 (2012).

9. Babalola, A.M. Visualization of Voxel Volume Emission and Absorption of Light in Medical Biology. Current Trends on Biostatistics and Biometrics 1, 71–81 (2019).

10. Tukora, B. Effective volume rendering on mobile and standalone VR headsets by means of a hybrid method. Pollack Periodica 15, 3–12 (2020).

11. Ruijters, D. & Vilanova, A. Optimizing GPU volume rendering. (2006).

12. Kruger, J. & Westermann, R. in IEEE Visualization, 2003. VIS 2003. 287-292 (2003).

13. Susaki, E.A. et al. Versatile whole-organ/body staining and imaging based on electrolyte-gel properties of biological tissues. Nat Commun 11, 1982 (2020).

14. Tainaka, K. et al. Chemical Landscape for Tissue Clearing Based on Hydrophilic Reagents. Cell Rep 24, 2196–2210 e2199 (2018).

15. Anderson, R.H., Spicer, D.E., Brown, N.A. & Mohun, T.J. The development of septation in the four-chambered heart. Anat Rec (Hoboken) 297, 1414–1429 (2014).

16. Captur, G. et al. Morphogenesis of myocardial trabeculae in the mouse embryo. J Anat 229, 314–325 (2016).

17. Anderson, R.H., Spicer, D.E., Mohun, T.J., Hikspoors, J. & Lamers, W.H. Remodeling of the Embryonic Interventricular Communication in Regard to the Description and Classification of Ventricular Septal Defects. Anat Rec (Hoboken) 302, 19–31 (2019).

18. Anderson, R.H., Brown, N.A. & Webb, S. Development and structure of the atrial septum. Heart 88, 104–110 (2002).

19. Garcia-Frigola, C., Shi, Y. & Evans, S.M. Expression of the hyperpolarization-activated cyclic nucleotide-gated cation channel HCN4 during mouse heart development. Gene Expr Patterns 3, 777–783 (2003).

20. Stieber, J. et al. The hyperpolarization-activated channel HCN4 is required for the generation of pacemaker action potentials in the embryonic heart. Proc Natl Acad Sci U S A 100, 15235–15240 (2003).

21. Wiese, C. et al. Formation of the sinus node head and differentiation of sinus node myocardium are independently regulated by Tbx18 and Tbx3. Circ Res 104, 388–397 (2009).

22. Hiruma, T., Nakajima, Y. & Nakamura, H. Development of pharyngeal arch arteries in early mouse embryo. J Anat 201, 15–29 (2002).

23. Yashiro, K., Shiratori, H. & Hamada, H. Haemodynamics determined by a genetic programme govern asymmetric development of the aortic arch. Nature 450, 285–288 (2007).

24. Kervrann, C., Legland, D. & Pardini, L. Robust incremental compensation of the light attenuation with depth in 3D fluorescence microscopy. J Microsc 214, 297–314 (2004).

25. Lloyd, S. Least squares quantization in PCM. IEEE Transactions on Information Theory 28, 129–137 (1982).

26. Arthur, D. & Vassilvitskii, S. in Proceedings of the eighteenth annual ACM-SIAM symposium on Discrete algorithms 1027–1035 (Society for Industrial and Applied Mathematics, New Orleans, Louisiana; 2007).

27. Wang, Z., Bovik, A.C., Sheikh, H.R. & Simoncelli, E.P. Image quality assessment: from error visibility to structural similarity. IEEE Trans Image Process 13, 600–612 (2004).

28. Bamforth, S.D. et al. Clarification of the identity of the mammalian fifth pharyngeal arch artery. Clin Anat 26, 173–182 (2013).

29. Rana, M.S., Sizarov, A., Christoffels, V.M. & Moorman, A.F. Development of the human aortic arch system captured in an interactive three-dimensional reference model. Am J Med Genet A 164A, 1372–1383 (2014).

30. Anderson, R.H., Bamforth, S.D. & Gupta, S.K. How best to describe the pharyngeal arch arteries when the fifth arch does not exist? Cardiol Young 30, 1708–1710 (2020).

31. Gupta, S.K., Gulati, G.S. & Anderson, R.H. Clarifying the anatomy of the fifth arch artery. Ann Pediatr Cardiol 9, 62–67 (2016).

32. Li-Villarreal, N., Rasmussen, T.L., Christiansen, A.E., Dickinson, M.E. & Hsu, C.W. Three-dimensional microCT imaging of mouse heart development from early post-implantation to late fetal stages. Mamm Genome 34, 156–165 (2023).

33. Ramirez, A. & Astrof, S. Visualization and Analysis of Pharyngeal Arch Arteries using Whole-mount Immunohistochemistry and 3D Reconstruction. J Vis Exp (2020).

