## Supplementary Figures for "Acto3D: user- and budget-friendly software for multichannel high-resolution three-dimensional imaging"

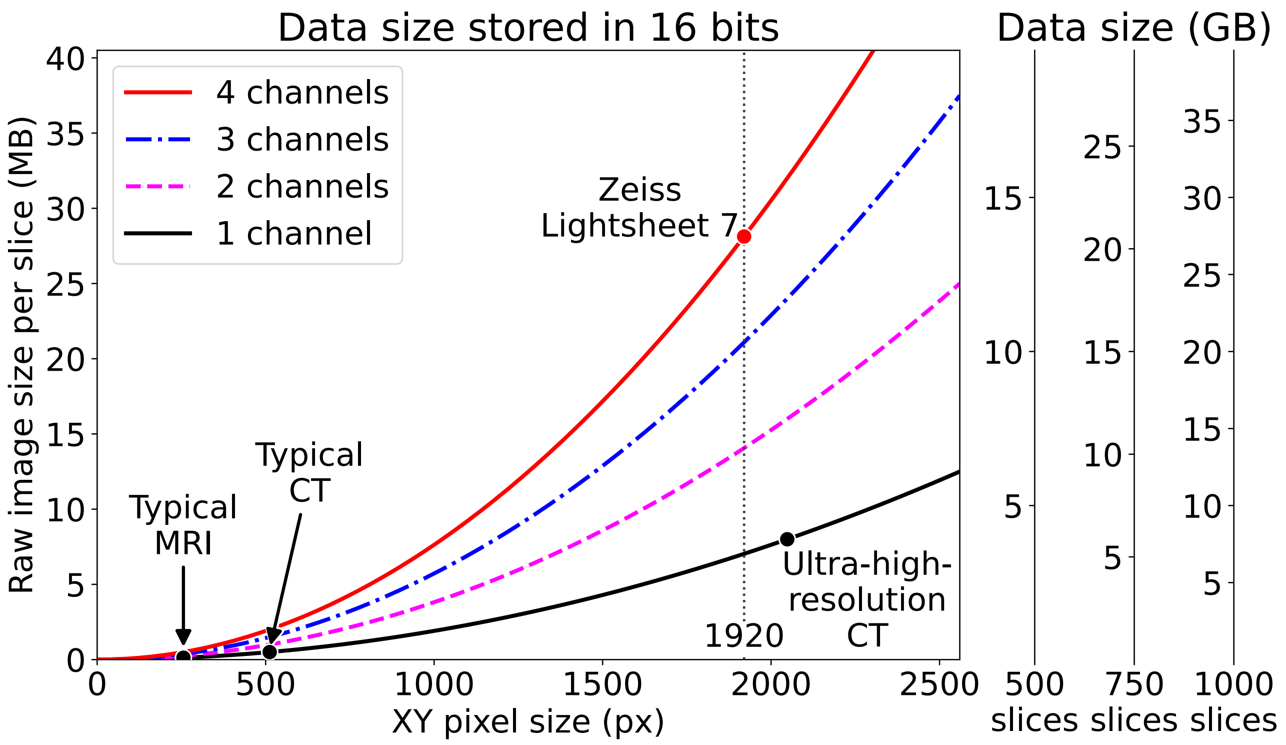


**Extended Data Fig. 1. Comparison of data size between medical imaging and fluorescence microscopy.** Data size per slice is compared for typical MRI (256 by 256 px), typical CT (512 by 512 px), ultra–high-resolution CT (2048 by 2048 px), and fluorescence microscopy (for example, 1920 by 1920 px for four-channel imaging with Zeiss Lightsheet 7). The right axis indicates the image data size when 500, 750, or 1000 slices are imaged.

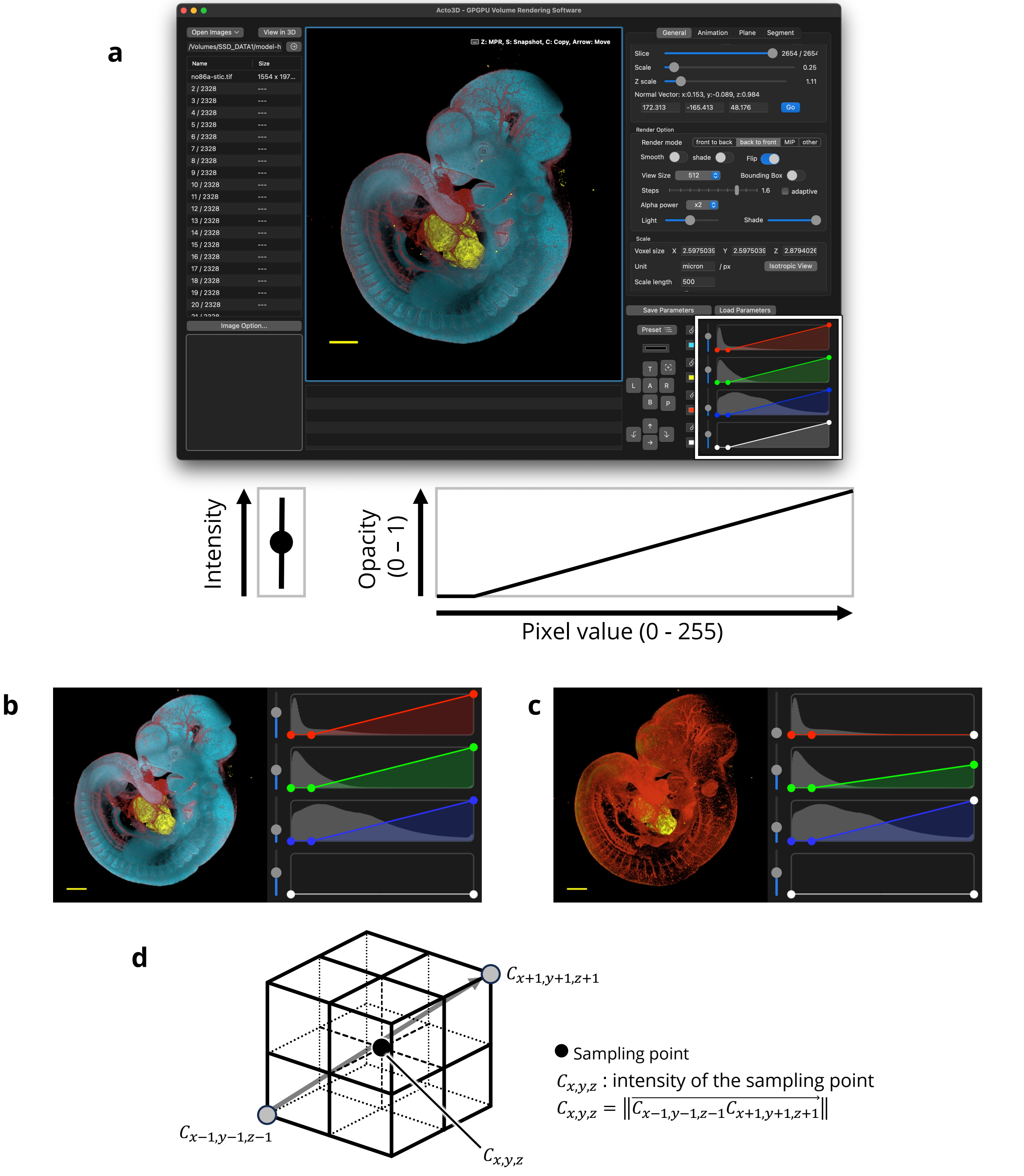


**Extended Data Fig. 2. Adjustment of visualization.**
**a**, Acto3D provides a panel within the software screen that allows the user to adjust color tones and the transfer function (white box). In addition to color tone, the user is thus able to adjust the intensity of each channel and the opacity corresponding to pixel values. The intensity setting multiplies the pixel value at each sampling point in each channel, so that increasing it makes the image brighter. Opacity is lowered for regions with low pixel values and set higher for those with high pixel values. In volume rendering, even very low opacities can make it difficult to see through the object if overlaid in large amounts; opacity is therefore squared and internally adjusted by default (this can be changed). The  mouse embryo showcased here corresponds to the stained embryos depicted in Figure 3. **b**, By increasing the intensity of the nuclear staining image, the surface of the entire embryo can be viewed. **c**, By reducing the intensity and opacity of the nuclear staining image, the internal structure can be viewed. **d**, For emphasis of edges, the norm of the difference in pixel value of surrounding points is used instead of the pixel value at the sampling point. It becomes possible to use this flexible transfer function by construction of a custom shader.

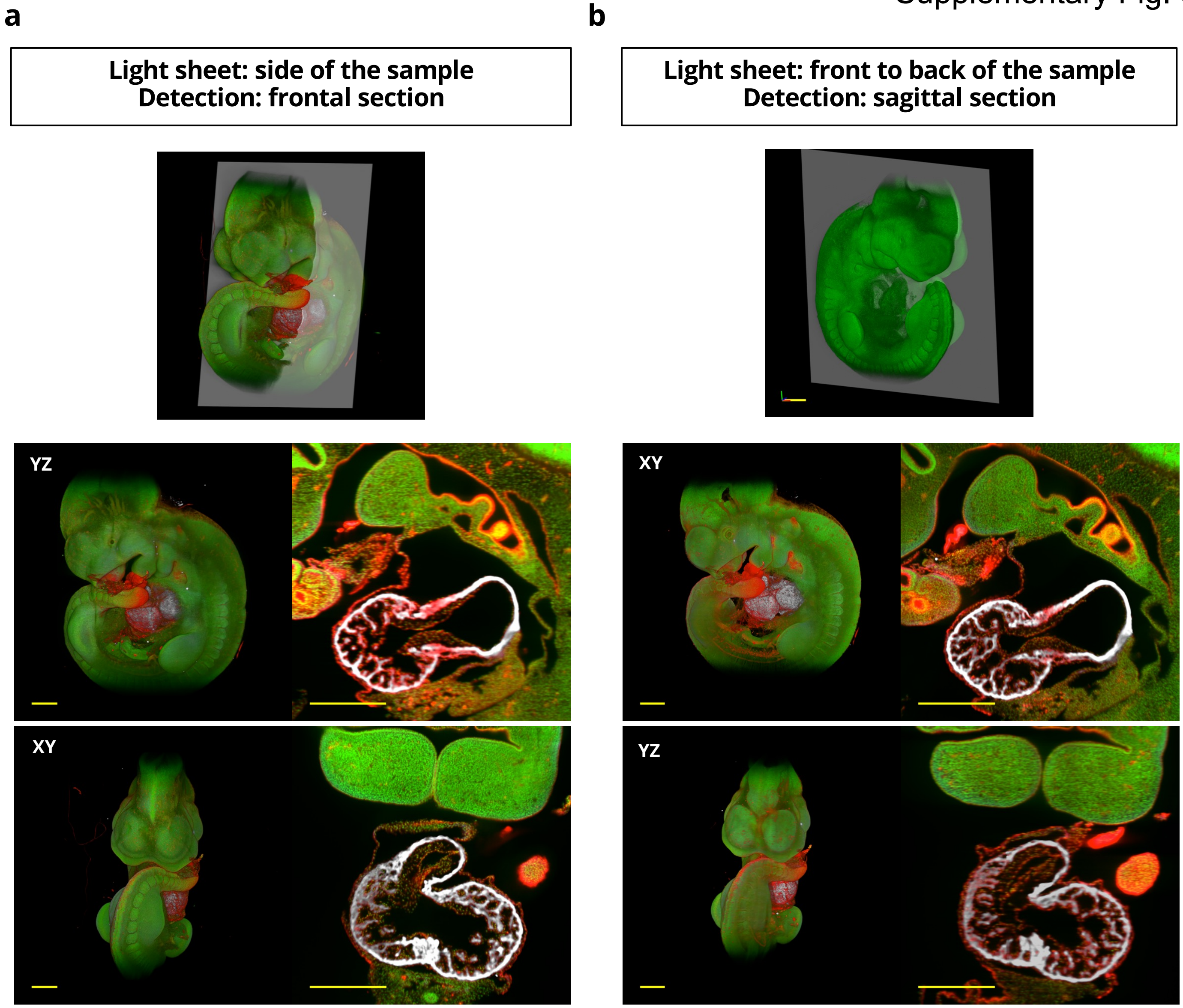


**Extended Data Fig. 3. Construction and visualization of 3D structures and MPR images from different illumination directions and sample orientations.**

For 3D reconstructions, Acto3D eliminates the need to precisely arrange the specimen and to align the light sheet plane in order to achieve maximum resolution. Comparison of data obtained from the same E10.0 mouse embryo stained as in Figure 3 by anteroposterior scanning in the coronal plane (**a**) with those obtained by lateral scanning in the sagittal plane (**b**) allowed the acquisition of XZ or YZ plane images that were similar in resolution to the theoretically highest-resolution plane image in the XY plane. MPR images are shown in the right panels of **a** and **b**. All scale bars, 500 µm.

Supplementary Fig. 4

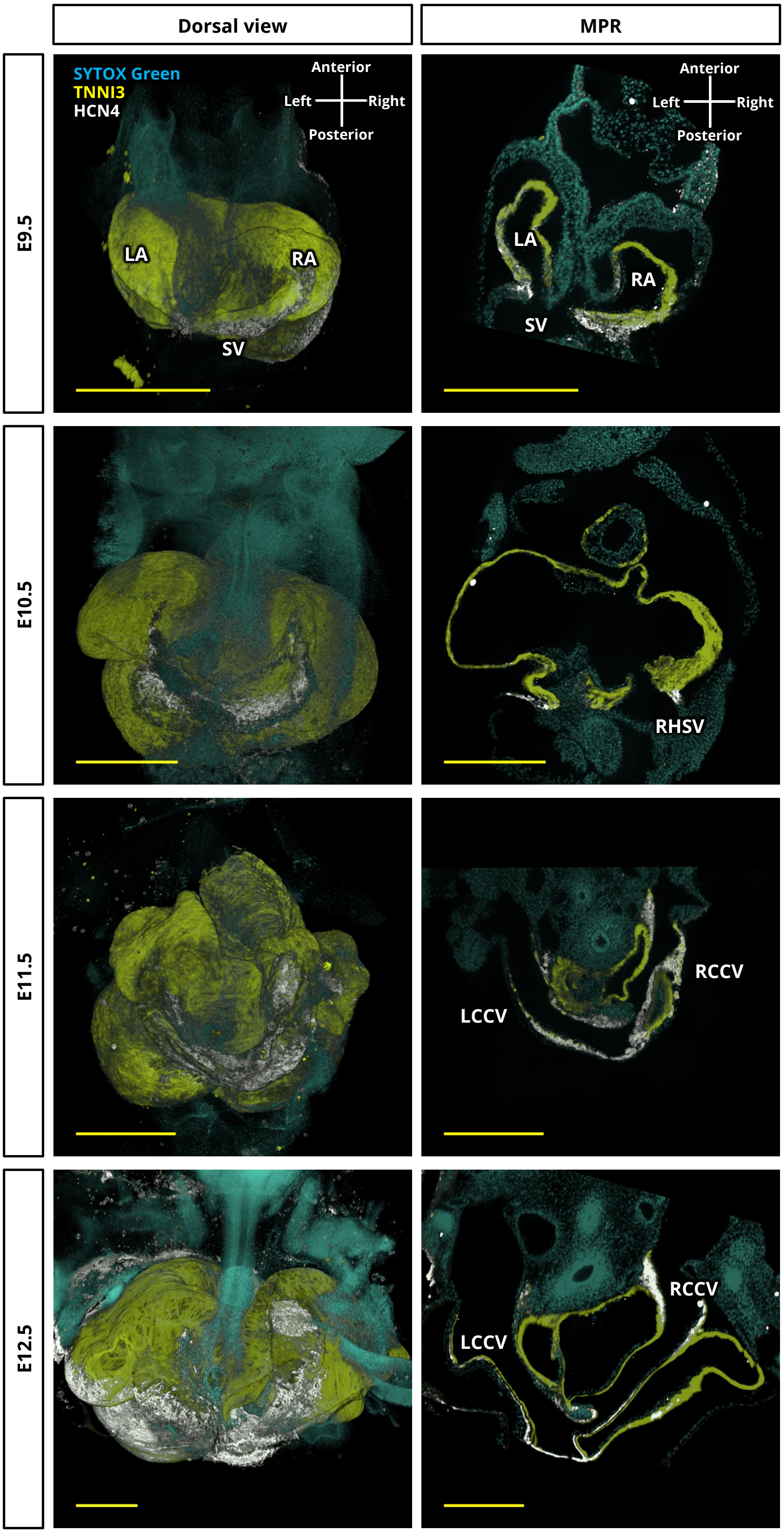


**Extended Data Fig. 4. 3D observation of HCN4 expression.**
The images depict views from the dorsal side of the heart of mouse embryos at E9.5, E10.5, E11.5, and E12.5, including MPR images from the same viewport showing the sinus venosus (SV) and common cardinal veins (CCVs). The embryos were stained with SYTOX Green and with antibodies to HCN4 and to TNNI3. The HCN4 signal was initially localized near the SV on the ventral side of the heart and gradually advanced upward toward the CCVs. The expression of HCN4 became left-right asymmetric in association with the left-right asymmetric remodeling of the cardinal veins. HCN4 expression overlapped with that of TNNI3. All scale bars, 500 µm. LA, left atrium; RA, right atrium; RHSV, right horn of SV; LCCV, left common cardinal vein; RCCV, right common cardinal vein.

Supplementary Fig. 5

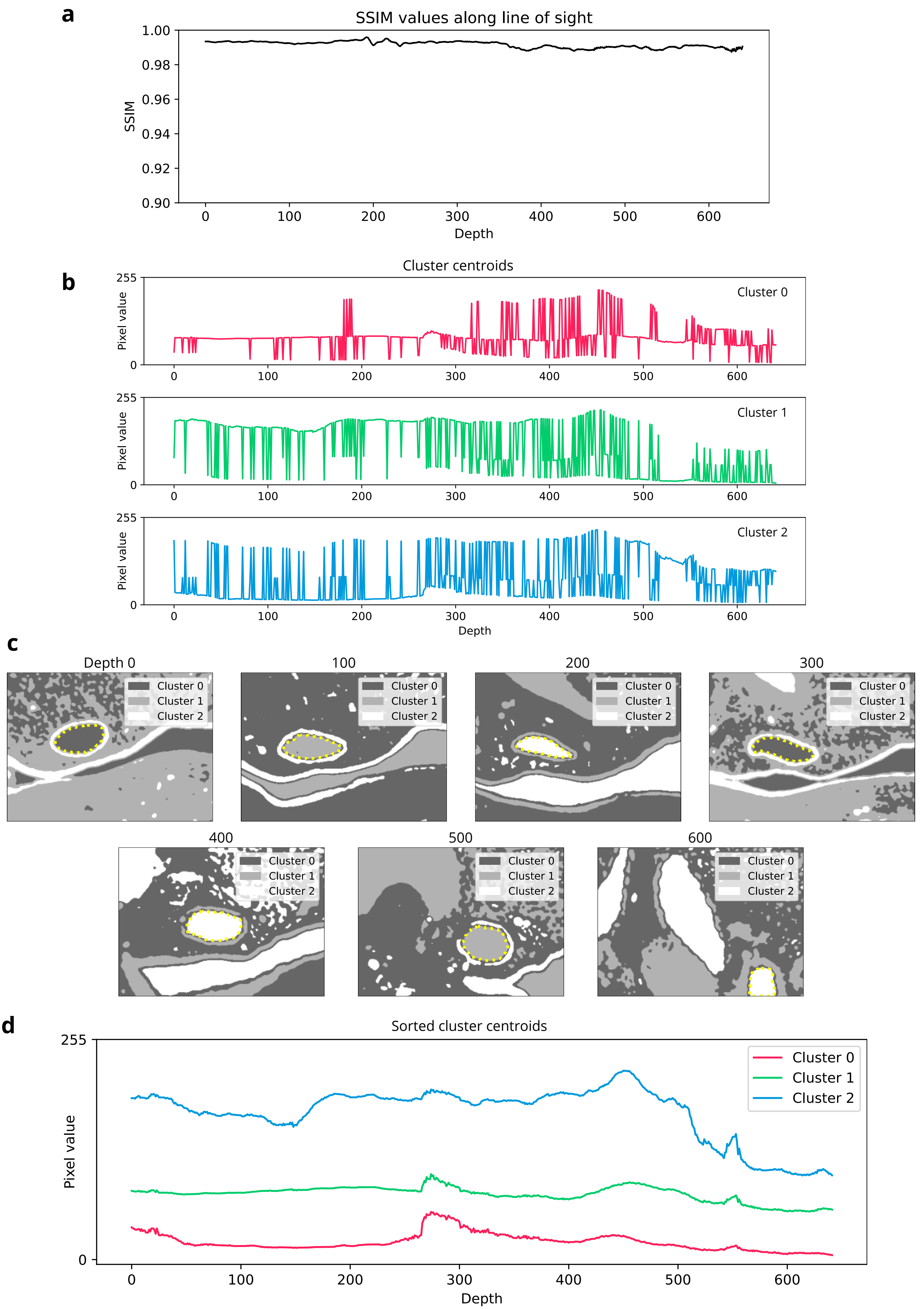

e

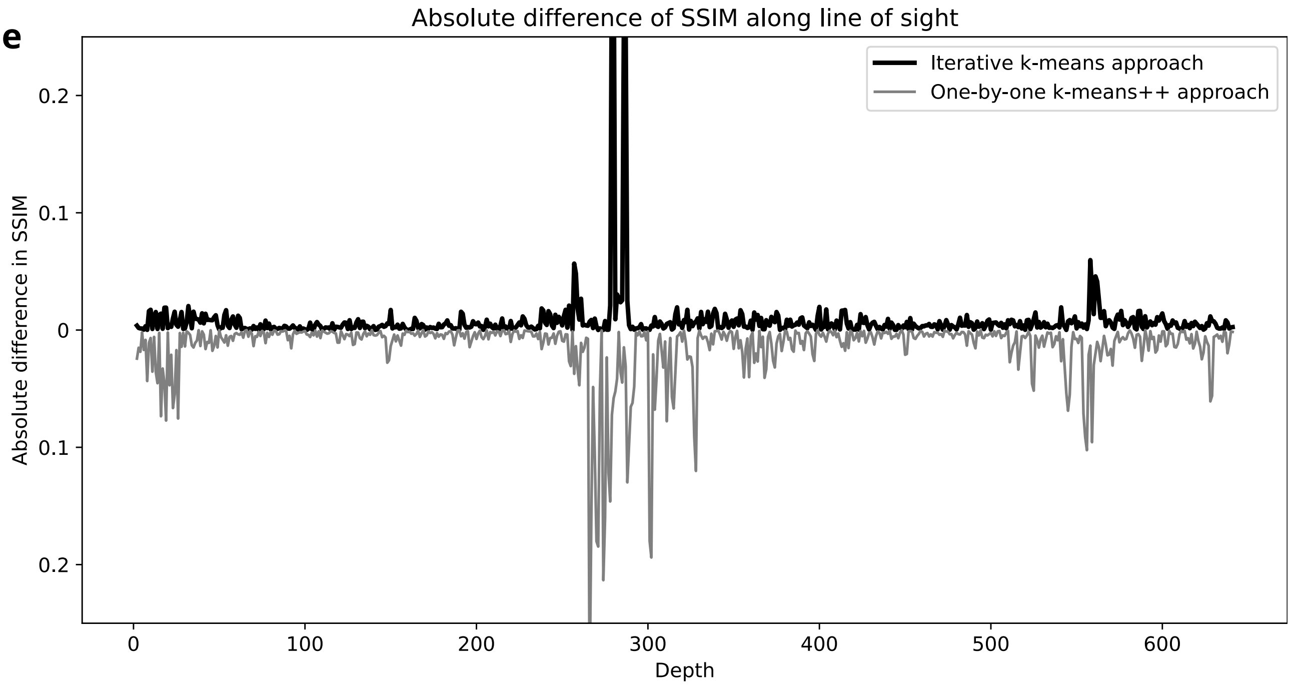


**Extended Data Fig. 5. Evaluation of the efficiency of iterative and one-by-one k-means++ approaches.** **a**, The similarity between adjacent slices along the line-of-sight direction is plotted as the Structural Similarity Index Measure (SSIM), which reflects similarity in structure and intensity. The high SSIM values of >0.98 indicate similarity in intensity and structure between adjacent slices. **b**, If k-means++ is applied independently to each cross-section—for instance, by setting the total number of clusters to 3—random numbers are assigned to each cluster in each cross-section. Note that cluster centroids of high, medium, and low pixel values are not consistently allocated to each cluster. **c**, When k-means++ is applied independently to each cross-section, clusters corresponding to the vascular lumen are assigned random numbers for each cross-section. **d**, The computed k-means++ clusters identified for each cross-section one-by-one in **b** were sorted to match those that should be classified into the same cluster on the image. Application of k-means++ independently to each cross-section one-by-one therefore entails additional computational costs. Note the lack of smoothness in the graph representing the pixel values of the cluster centroids in comparison with the results obtained by the iterative k-means++ method (Fig. 5b). **e**, The clustered grayscale images obtained by the iterative (Fig. 5c) or one-by-one (**c**) k-means++ approaches were subjected to SSIM calculation. The absolute difference in SSIM was computed as $|\mathrm{SSIM}_{n+1} - \mathrm{SSIM}_n|$, where $\mathrm{SSIM}_n$ denote the SSIM values calculated from the $n$th and $(n+1)$th clustered images, respectively. The results reveal that neighboring images are more similar with the iterative method (with the absolute difference in SSIM being close to 0) compared with the one-by-one k-means++ approach.

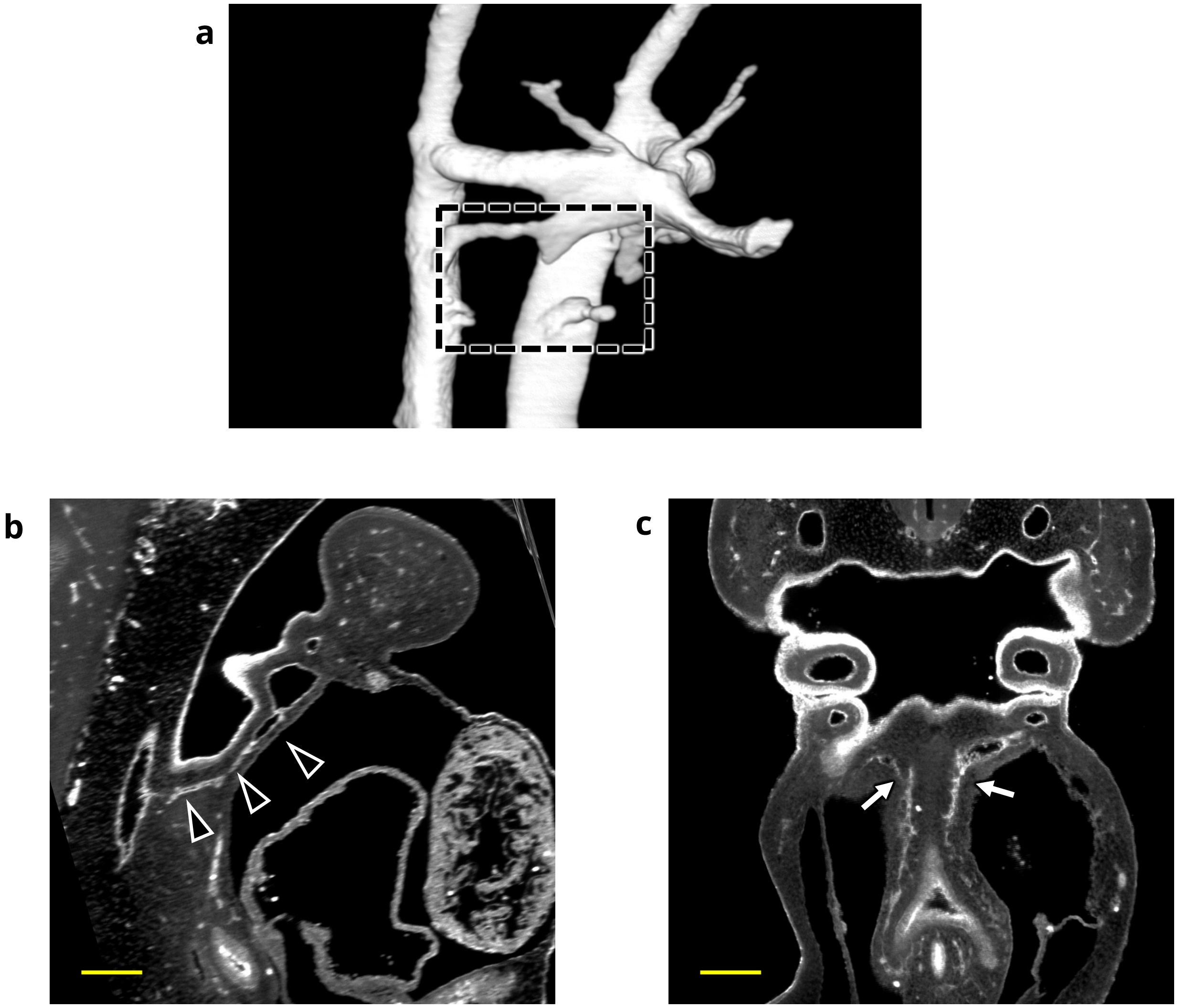


**Extended Data Fig. 6. Issues in constructing 3D models of extremely narrow blood vessels.**
**a**, A 3D reconstructed image of PAAs in a mouse embryo at E10.0 (32-somite stage). The specimen is the same as that in Figure 6e. **b**,**c**, MPR views of the same embryo as in **a** are shown as sagittal (**b**) and frontal (**c**) sections. In the 3D model (**a**), the sixth PAA did not appear continuous between the aortic sac and dorsal aorta. However, in the MPR views of the region corresponding to that where the sixth PAA was expected to be identified in **a** (dashed black square), a continuous line of endothelium (black arrowheads with a white border) was observed between the aortic sac and dorsal aorta, suggesting that the artery was not constructed as a result of it being too thin or not possessing a lumen. The endothelial structure of the pulmonary artery (white arrow with a black border) was observed, but due to its extreme thinness, it could not be constructed. Scale bars, 200 μm.
